# Essential Role of Protein Kinase R in the Pathogenesis of Pulmonary Veno-occlusive Disease

**DOI:** 10.1101/2025.04.18.649514

**Authors:** Amit Prabhakar, Rahul Kumar, Meetu Wadhwa, Abhilash Barpanda, Joseph Lyons, Asavari Gowda, Simren Gupta, Ananyaa Arvind, Prajakta Ghatpande, Arun P. Wiita, Brian B. Graham, Giorgio Lagna, Akiko Hata

## Abstract

Pulmonary veno-occlusive disease (PVOD) is a rare and severe subtype of pulmonary arterial hypertension, marked by progressive remodeling of small pulmonary arteries and veins with no therapies. Using a mitomycin C (MMC)-induced rat model, we previously demonstrated that protein kinase R (PKR)-mediated integrated stress response (ISR) drives endothelial dysfunction and vascular remodeling. To determine if PKR is the sole mediator of ISR and the pathogenesis, we treated control (Ctrl) and PKR knockout (KO) mice with the same dose of MMC. Consistent with rat data, Ctrl mice displayed ISR activation, vascular remodeling, and pulmonary hypertension after MMC treatment, while KO mice showed none of these phenotypes. Proteomic analysis revealed that MMC-mediated ISR activation attenuates protein synthesis in Ctrl but not in KO mice. These findings underscore the significance of PKR-dependent ISR activation and subsequent perturbation of proteostasis as central mechanisms driving PVOD pathogenesis and identifying PKR as a promising therapeutic target.

## Introduction

Pulmonary veno-occlusive disease (PVOD) is a severe form of pulmonary arterial hypertension (PAH), characterized by progressive remodeling and obstruction of small pulmonary arteries (PAs), veins (PVs), and capillaries (1–3). The five-year survival rate for PVOD patients is 27%(4), significantly lower than 61% for PAH patients (5). PVOD affects both sexes equally and is estimated to occur in 1 in 5-10 million people per year (1–3). Due to the similarities in radiographic findings between PVOD and PAH, PVOD patients are frequently misdiagnosed as having PAH(2, 3). Typically, patients with PVOD respond poorly to PAH therapies(2, 3). Furthermore, when treated with PAH-specific drugs, PVOD patients can develop life-threatening pulmonary edema(2, 3). Thus, there is a critical need for therapeutics specifically tailored to PVOD.

Biallelic mutations in the *Eif2ak4* gene are the leading genetic cause of PVOD(6). *Eif2ak4* encodes general control nonderepressible 2 (GCN2, also known as EIF2AK4), one of the four eIF2 kinases, along with heme-regulated inhibitor (HRI, also known as EIF2AK1), protein kinase R (PKR, or EIF2AK2), and PKR-like endoplasmic reticulum kinase (PERK, also known as EIF2AK3), that become active in response to different cellular stresses and phosphorylate serine-51 (S51) of the alpha subunit of eukaryotic initiation factor 2 (eIF2α)(7). Upon phosphorylation of eIF2α, global translation is attenuated, thereby conserving energy, reprogramming gene expression, and restoring proteostasis, a physiological response to stress collectively known as the integrated stress response (ISR) (8, 9). The ISR is essential for an organism’s adaptation to stress conditions and changing environments (8, 9). When cap-dependent translation is globally suppressed, the transcripts with upstream open reading frames, such as cyclic AMP-dependent transcription factor 4 (ATF4) and growth arrest and DNA damage-inducible protein 34 (GADD34; also known as PPP1R15A), are preferentially translated (8, 9). Upon activation of the ISR, the basic-leucine zipper transcription factor ATF4 accumulates and translocates to the nucleus, where it binds to stress response genes (SRGs) and activates the transcription. The induction of SRGs facilitates organisms to manage stress, maintain fitness, and adapt to new environmental conditions (9).

In addition to the *Eif2ak4* mutations, exposure to alkylating agents, such as mitomycin C (MMC), cyclophosphamide, and cisplatin, is known to induce PVOD in cancer patients (10–12). Indeed, the administration of MMC to rats causes a spectrum of PVOD phenotypes, including right ventricular hypertrophy and pulmonary vascular lesions such as medial thickening, luminal obstruction in PAs and PVs, intimal obliteration, adventitial growth, and thrombosis. These features make it a valuable animal model for studying PVOD (11, 13–17). Using the MMC-induced PVOD model in rats, we demonstrated that MMC induces the activation of one of the eIF2 kinase PKR, which subsequently triggers the ISR(16, 17). This leads to the depletion of the vascular endothelial adhesion molecule vascular endothelial-cadherin (VE-Cad) in complex with Rad51, contributing to increased permeability and vascular remodeling (16). Administration of a small molecule inhibitor of PKR, C16, or ISR, ISRIB prevents and attenuates ISR activation and PVOD pathogenesis in rats(16, 17). PKR regulates the innate immune response to viral infections (18). It is activated by binding to double-stranded RNA (dsRNA) introduced during viral infection and replication(19). Several stimuli, including DNA damage, pro-inflammatory cytokines, and heat shock, are also known to activate PKR(19). The mechanism of PKR activation by MMC and the cell-type specific responses downstream of the PKR-ISR axis remains unclear.

The potential involvement of the other three eIF2 kinases in activating the ISR and contributing to the development of cardiovascular phenotypes of PVOD in response to MMC remains unclear, given that all four eIF2 kinases are expressed in pulmonary vascular endothelial cells (PVECs) (16). In this study, we administered MMC to PKR homozygous null mice and examined the cardiopulmonary phenotypes associated with PVOD (20). Upon MMC treatment, wild-type littermates showed signs of active ISR and developed PVOD-like phenotypes as early as five days post-treatment. In contrast, PKR-deficient mice do not exhibit signs of ISR activation or PVOD phenotypes after MMC treatment. This study reports the first murine model of PVOD and demonstrates that PKR is an essential mediator of ISR and the onset of PVOD-like vascular remodeling.

## Results

### No ISR activation following MMC treatment in PKR-deficient mice

To study the role of PKR in the MMC-mediated pathogenesis of PVOD, 9-10 weeks old homozygous PKR knockout (KO) in the C57Bl/6 background mice in which both copies of the *Eif2ak2* gene that encodes PKR were ablated (*Eif2ak2 −/−*)(20, 21) and the littermate control (Ctrl) mice (*Eif2ak2 +/+*) were administered with PBS (vehicle; Veh) or MMC. KO mice develop and breed with no phenotypes and have an average lifespan (20, 21). Five days after the administration of Veh or MMC, total lung lysates were subjected to the immunoblot to examine the activation status of PKR and ISR. The levels of active PKR were reviewed as the amount of phosphorylated-PKR (p-PKR) auto-phosphorylated at threonine-446 (p-PKR) relative to total PKR (t-PKR) (p-PKR/t-PKR ratio) and found that it was increased 2.6-fold in MMC-treated Ctrl mice relative to Veh-treated Ctrl mice (**Fig. 1A**), suggesting MMC-induces PKR activation in Ctrl mice like in rats(16, 17). Furthermore, the amount of the S51-phosphorylated eIF2α (p-eIF2α) relative to total eIF2α (t-eIF2α) (p-eIF2α/t-eIF2α ratio) and ATF4 were increased 1.3-fold and 4.3-fold in Ctrl mice following MMC treatment, respectively (**Fig. 1A**). These results in Ctrl mice demonstrate MMC mediates the activation of PKR and the ISR, which involves eIF2α phosphorylation and inhibition of cap-dependent translation, and preferential translation of ATF4 mRNA, similar to findings in rats (16, 17). As expected, p-PKR and t-PKR were below detectable levels in KO mice, validating the genetic ablation of PKR in KO mice (**Fig. 1A**). As observed in rats (17), MMC treatment reduced the amount of the regulatory subunit of protein phosphatase 1 (PP1) GADD34, which is essential for the dephosphorylation of eIF2α and inactivation of the ISR (**Fig. 1A**), indicating that MMC mediates constitutive eIF2α phosphorylation and ISR activation in Ctrl mice. The levels of mRNA (**Supple. Fig. 1A**) and protein (**Supple. Fig. 1B**) for other eIF2 kinases, such as GCN2, PERK, and HRI, were comparable between Ctrl and KO mice regardless of the treatments. In contrast with Ctrl mice, KO mice showed no significant change in the p-eIF2α/t-eIF2α ratio or ATF4 protein levels following MMC treatment, suggesting an absence of ISR activation in PKR-deficient mice (**Fig. 1A**).

**Fig. 1.**
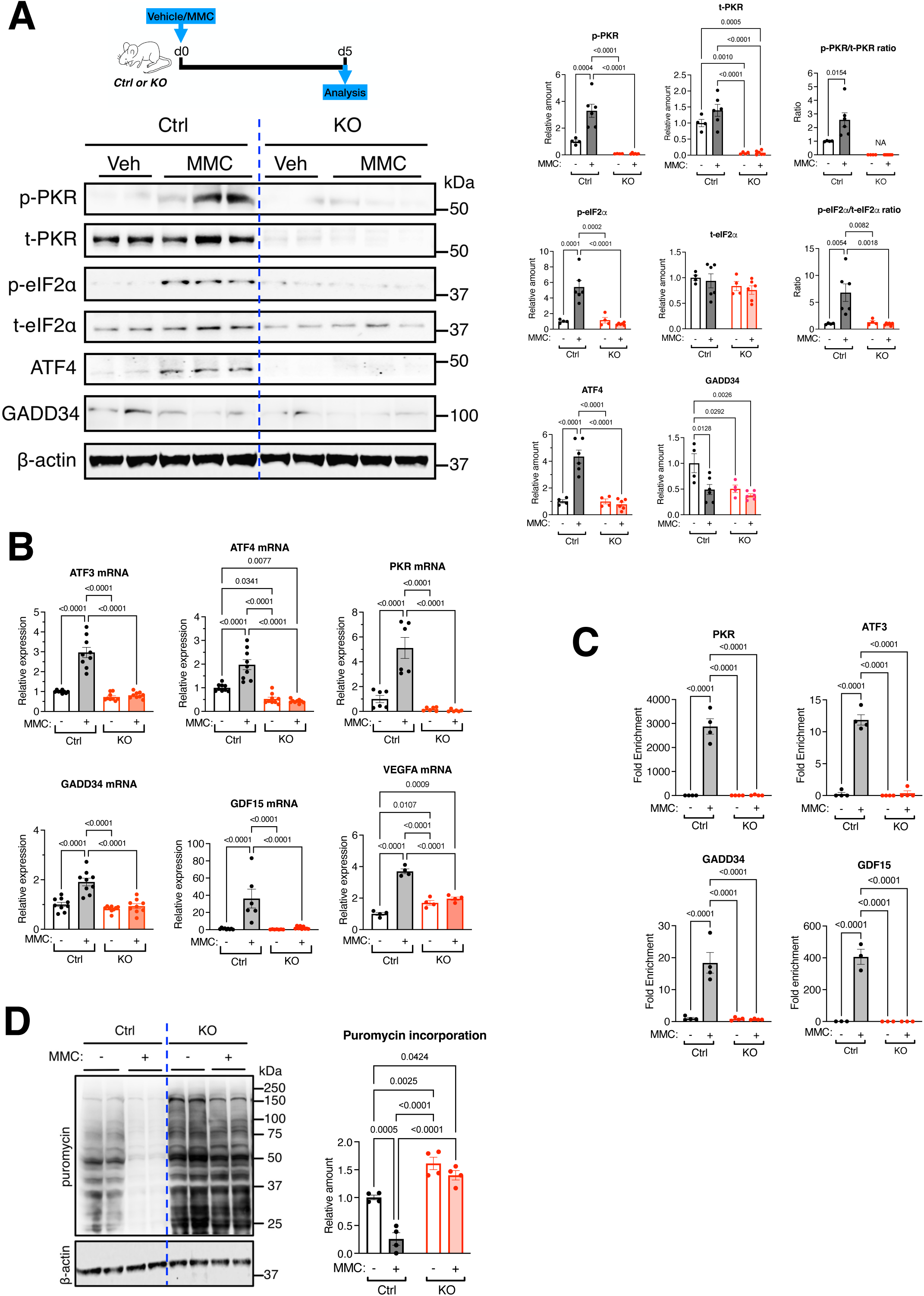
ISR activation upon MMC treatment is abolished in PKR-deficient mice. **A.** Schematic representation of MMC-induced PVOD mouse model (top) and immunoblot analysis of the indicated proteins in total lung lysates from vehicle (Veh) or MMC-treated Control (Ctrl) and PKR-KO (KO) mice on day 5 (bottom, left). The relative amounts of the indicated proteins, normalized to β-actin, are shown as mean ± SEM (bottom right). n=2-3 independent samples. **B.** The levels of ATF4 target gene mRNAs, such as *ATF3, ATF4, PKR, GADD34, GDF15,* and *VEGFA*, in the lungs of Ctrl and KO mice treated with Veh or MMC on day 5 were analyzed by qRT-PCR. The results were normalized to β-actin levels and are presented as mean ± SEM. n=4-6 independent samples. **C.** ChIP assay was performed using the lungs harvested from Ctrl and KO mice treated with Veh or MMC employing an anti-ATF4 antibody, followed by PCR amplification corresponding to the genomic region of the PKR, ATF3, GADD34, and GDF15 gene spanning the ATF4 binding sequence. The PCR results are presented as fold enrichment over the input as mean ± SEM. n=4 independent experiments. **D.** In vivo puromycin incorporation assay. Ctrl and KO mice treated with Veh or MMC were administered puromycin on day 5, followed by lung lysate preparation. After normalizing the total protein content, samples were subjected to SDS-PAGE. Puromycin-labeled proteins were visualized by immunoblotting using anti-puromycin and anti-β-actin antibodies (left panel). The levels of puromycin-labeled proteins, normalized to β-actin as a loading control, are shown as mean ± SEM (right panel). n=4 independent samples per group.

As previously observed in rats(16, 17), the levels of transcripts of the ATF4 target genes, such as *ATF3, ATF4, PKR, GADD34, growth differentiation factor 15 (GDF15), and vascular endothelial growth factor A (VEGFA)* (*22–25*), were increased 3.0-fold, 2.0-fold, 8.8-fold, 1.9-fold, 36.1-fold, and 3.7-fold, respectively, following MMC treatment in Ctrl mice (**Fig. 1B**). Despite the increase in GADD34 mRNA levels (**Fig. 1B**), GADD34 protein levels decreased following MMC treatment in Ctrl mice (**Fig. 1A**), similar to findings in rats (16, 17). This suggests that post-transcriptional regulation of GADD34 leads to the reduction of GADD34 protein levels after MMC treatment. In contrast with Ctrl mice, no induction of ATF4 target gene transcripts was observed in KO mice following MMC treatment (**Fig. 1B**). Mice heterozygous for PKR (Eif2ak2 +/-) displayed a reduced yet significant induction of ATF4 target gene transcripts, suggesting that PKR haploinsufficiency leads to a partial induction of ATF4 target genes (**Supple. Fig. 2**). The chromatin immunoprecipitation sequencing (ChIP-seq) experiment for ATF4 by the ENCODE consortium revealed the genome-wide ATF4 binding sites (BS) in human lymphoblast K562 cells (Accession No. ENCFF484GNY and ENCFF742FPU), which include the PKR, ATF3, GADD34, and GDF15 genes (**Supple. Fig. 3**). After the administration of Veh or MMC, we performed a ChIP assay using the total lung lysates harvested from Ctrl mice and confirmed that the genomic fragments of *PKR*, *ATF3*, *GADD34* and *GDF15* containing the ATF4 BS (**Supple. Fig. 3**) were enriched 2,875-fold, 12-fold, 18-fold, and 355-fold, respectively, in MMC-treated Ctr mice over Veh-treated Ctrl mice (**Fig. 1C, bottom**). However, we did not detect an association of ATF4 with any of these genes following MMC treatment in KO mice (**Fig. 1C**), which is consistent with no induction of the PKR, ATF3, and GADD34 transcripts in KO mice upon MMC treatment (**Fig. 1B**). These results demonstrate that PKR is essential for the MMC-mediated induction of ATF4 and the transcriptional activation of ATF4 target genes. The in vivo puromycin incorporation assay revealed a 74% reduction in nascent protein synthesis in the lung of Ctrl mice following MMC treatment (**Fig. 1D**), validating the ISR activation (8, 9). In KO mice, however, there was no reduction in the nascent protein synthesis after MMC treatment, confirming no ISR activation upon MMC treatment in KO mice (**Fig. 1D**). We noted the levels of protein synthesis in Veh-treated KO mice were 1.6-fold higher than those in Veh-treated Ctrl mice, indicating that the deletion of PKR slightly increases the global protein synthesis (**Fig. 1D**). These results demonstrate that PKR is solely responsible for the MMC-mediated ISR activation and attenuation of global protein synthesis.

**Fig. 2.**
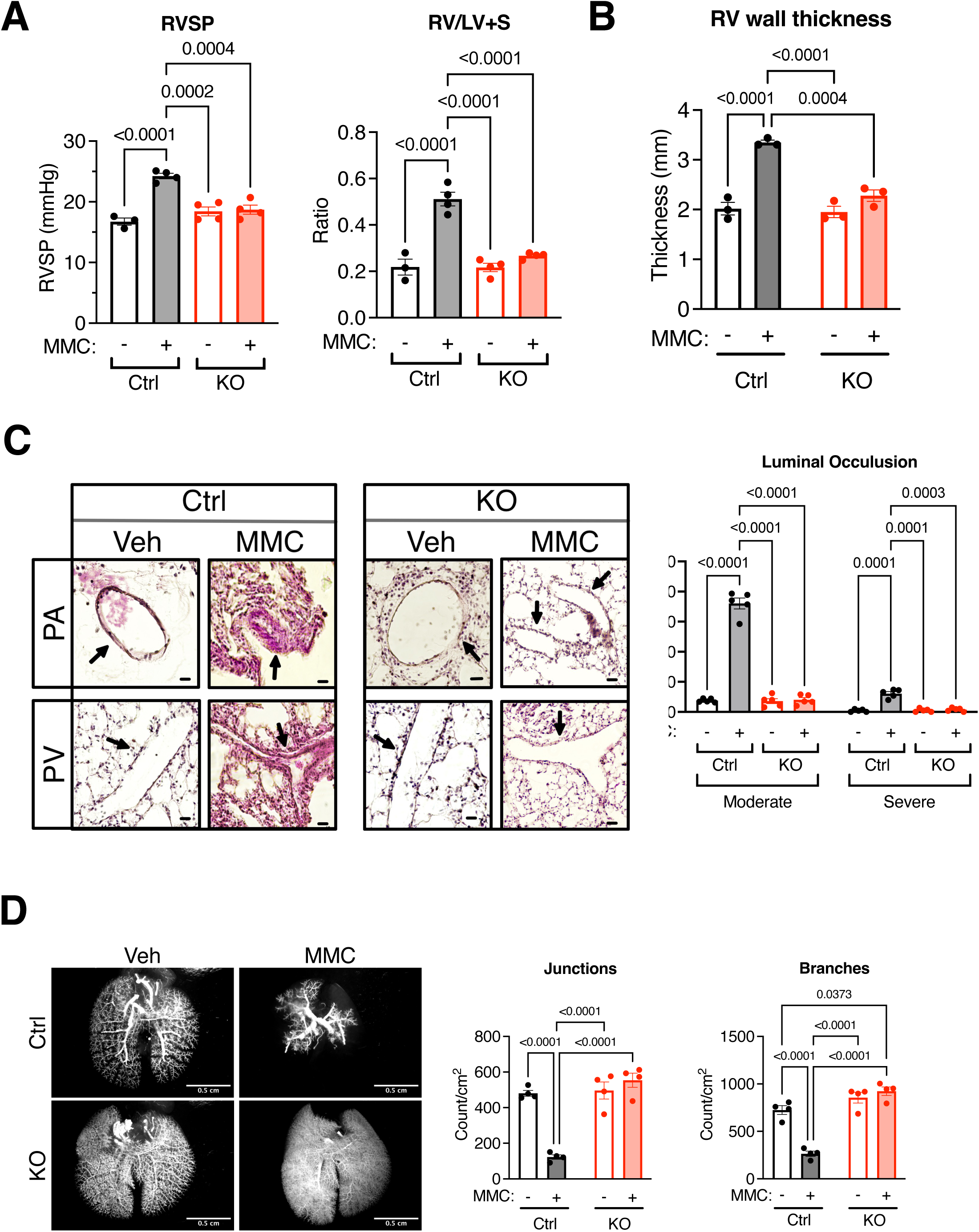
PKR-deficient mice do not develop PVOD phenotypes. **A.** RVSP (mmHg) (n=3-4 independent samples per group) and RV/LV+S ratio (n=3-4 independent samples per group) in the vehicle (Veh) or MMC-administered control (Ctrl) and KO mice. n=3-4 independent samples per group. **B.** RV wall thickness (mm) of Veh- or MMC-treated Ctrl and KO mice was measured and shown as mean+SEM (left). The wall thickness was measured at five locations, and the mean value was calculated per sample. n=3 independent samples per group. **C.** H&E staining of pulmonary vessels (PA and PV; arrows) in Ctrl and KO mice administered with Veh or MMC (left). The third column is a magnified image of the black rectangle area in the second column (left). The fraction (%) of moderately (25-40% and severe is above 40% occlusion) and severely (>40% occlusion) occluded vessels were counted and shown as mean+SEM (right). Scale bar=10 mm. n=5 independent samples. **D.** Microfil casting of the lung vasculature in Ctrl and KO mice treated with either Veh or MMC on day 5. Holistic images of the entire lung are displayed on the left, with a scale bar representing 0.5 cm. The number of junctions and branches per cm² of distal pulmonary vessels was quantified, with the data presented as mean ± SEM (right). n = 3 independent samples per group.

**Fig. 3.**
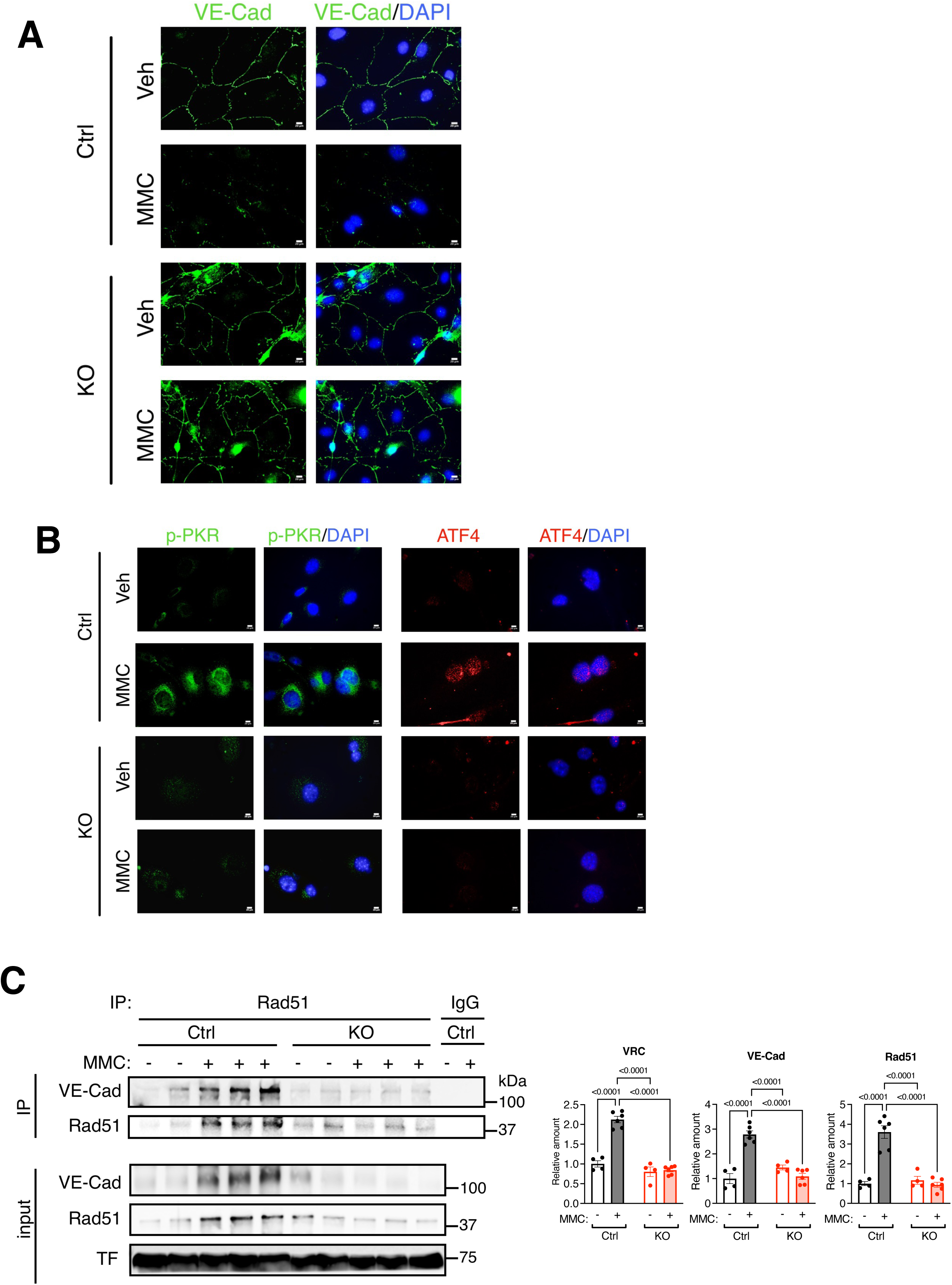
MMC treatment does not impair pulmonary vascular endothelium in PKR-deficient mice following MMC treatment. **A.** PVECs were isolated from the lungs of Ctrl and KO mice administered with vehicle or MMC and subjected to IF staining with anti-VE-Cad antibody (green) and DAPI (blue) for nuclei. Scale bar=10 mm. **B.** PVECs isolated from the mice administered with vehicle or MMC were subjected to IF staining with p-PKR (green, left), ATF4 (red, right), and DAPI (blue) for nuclei. Scale bar=10 mm. **C.** The plasma isolated from Veh or MMC-treated Ctrl and KO mice on day 5 were subjected to an IP with an anti-Rad51 antibody or nonspecific IgG (control), followed by immunoblot analysis of an anti-VE-Cad (for VRC) and anti-Rad51 antibody to detect the interaction between these proteins. Immunoblot with an anti-TF antibody (TF) is shown as loading control. n=3 independent samples (top). The relative amounts of the indicated proteins normalized to TF are demonstrated as mean ± SEM (bottom). *n* = 4-6 independent samples.

### PKR-deficient mice do not develop PVOD

To examine whether MMC mediates pulmonary hypertension in the absence of PKR, Ctrl and KO mice were administered with vehicle (Veh) or MMC. On day 5 after the MMC injection, we measured the RV systolic pressure (RVSP) for PA pressure and the ratio of the RV weight to the left ventricle (LV) plus septum (S) weight (RV/LV+S) for RV hypertrophy. Ctrl mice treated with MMC exhibited an elevation of RVSP (from 16.7±0.6 mmHg to 24.2±0.5 mmHg after five days; however, no increase in RVSP was detected in KO mice following MMC treatment (from 18.4±0.7 mmHg to 18.7±0.7 mmHg) (**Fig. 2A**). The basal and MMC-induced RVSP were comparable between male and female Ctrl and KO mice (**Supple. Fig. 4**), similar to findings observed in rats (16, 17). Following MMC treatment, the RV/LV+S ratio increased significantly in Ctrl mice, from 0.22±0.03 (Veh) to 0.51±0.03 (MMC) (**Fig. 2A**). In contrast, in KO mice, the RV/LV+S ratio remained not significantly post-MMC (0.27±0.00), with values comparable to Veh-treated mice (0.22±0.02) (**Fig. 2A**), indicating no development of RV hypertrophy in KO mice post-MMC. The deletion of PKR did not affect RVSP or the RV/LV+S ratio, as there were no significant differences observed in these parameters between Veh-treated Ctrl and KO mice (**Fig. 2A**). We also observed pleural and pericardial effusion, conditions associated with PVOD patients (2), in all MMC-treated Ctrl mice. However, none of the MMC-treated KO mice developed pleural or pericardial effusion. Fifty % of MMC-treated Ctrl mice died by day 8. In contrast, no mortality was observed in MMC-treated KO mice up to day 15 (**Supple. Fig. 5**), demonstrating the absence of MMC-induced cardiovascular phenotypes in KO mice. Ctrl mice, but not KO mice, showed a 66% increase in RV wall thickness following MMC treatment (**Fig. 2B** and **Supple. Fig. 6**), confirming the development of RV hypertrophy in MMC-treated Ctrl mice. These results demonstrate that PKR-null mice are protected from developing the phenotypes of PVOD upon MMC administration. H&E staining of the lung showed that Ctrl mice developed medial hyperplasia and hypertrophy in PAs and PVs following MMC treatment (11, 13) (**Fig. 2C, left**). The fraction of PAs and PVs with moderate (25-40% occlusion) and severe (more than 40% occlusion) occlusion in Ctrl mice increased 9.5-fold (from 3.8% to 36.0%) and 14-fold (from 0.4% to 6.1%), respectively, following MMC treatment (**Fig. 2C, right**). MMC-treated Ctrl mice that died on day 5 after MMC administration revealed severe remodeling of PAs and PVs that resulted in complete occlusion of the lumen (**Supple. Fig. 7**). KO mice did not develop vessel occlusion following MMC treatment (**Fig. 2C, right**), which is consistent with the lack of increase in the RVSP and RV/LV+S ratio in MMC-treated KO mice (**Fig. 2A**). There were no signs of spontaneous development of vascular remodeling in Veh-treated KO mice (**Fig. 2C, left**). Casting of vessels with Microfil revealed obstructions in the distal pulmonary vessels of MMC-treated Ctrl mice (**Fig. 2D, left**). Quantitative analysis demonstrated 74.4% and 63.5% fewer junctions and branches in the pulmonary vasculature of MMC-treated Ctrl mice compared to Veh-treated Ctrl mice, respectively (**Fig. 2D, right**). In contrast, MMC treatment did not change the number of junctions or branches in KO mice (**Fig. 2C, right**). The histological findings (**Fig. 2C**) confirm that MMC induces pulmonary vascular remodeling in mice, similar to the pathology observed in PVOD patients and previously documented in rats(16, 17). Consistent with findings from the rat model of PVOD (16, 17), no vascular remodeling was observed in other organs, including the brain, kidney, heart, liver, and intestine of Ctrl or KO mice following MMC treatment (**Supple. Fig. 8**), indicating that MMC-mediated vascular remodeling is specific to the lung. These results demonstrate that MMC triggers neither ISR activation nor pulmonary vascular remodeling without PKR, confirming that PKR is solely responsible for MMC-mediated PVOD. These findings also rule out the involvement of translational regulation mechanisms independent of eIF2 kinases or the ISR, such as mTOR complex 1(26).

**Fig. 4.**
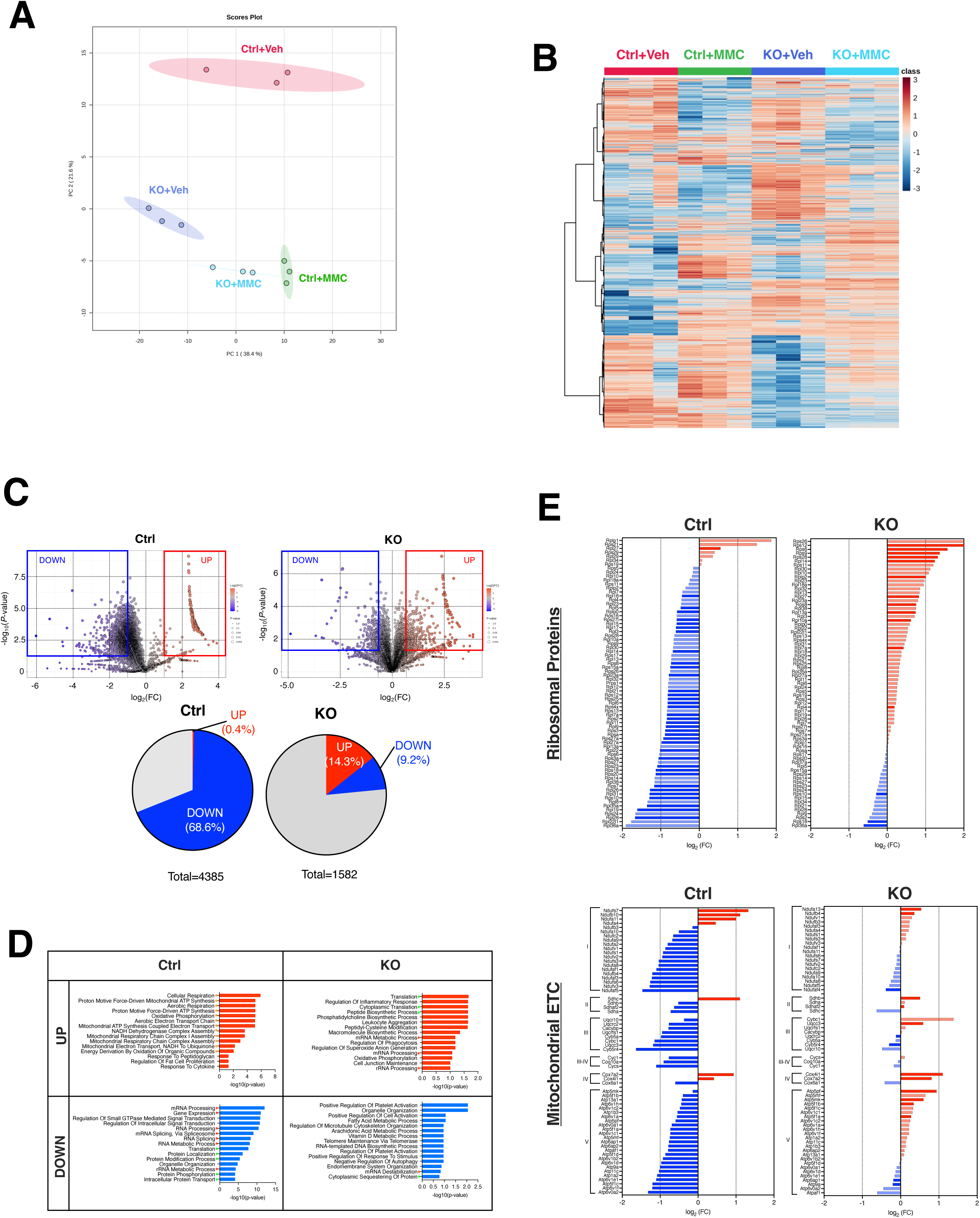
Effects of MMC on the lung proteomic landscape in Ctrl and KO mice. **A.** Lungs harvested on day 5 from Veh- or MMC-treated Ctrl and KO mice were subjected to MS analysis. The principal component analysis of the MS data for lungs from Veh- or MMC-treated Ctrl and KO mice, conducted in triplicate, is shown. **B.** A hierarchically clustered heat map of differentially expressed proteins (DEPs) that are statistically significant (*p* <0.05 by two-tailed Student’s t test) in the lungs of Veh- or MMC-treated Ctrl and KO mice. n=3 per group. **C.** Volcano plots compare the proteome of MMC-treated versus Veh-treated Ctrl and KO mouse lungs (top). A larger circle size represents a lower *P*-value, while the color gradient from blue to red corresponds to increasing log_2_(FC) values, with blue indicating lower values and red indicating higher values. Pie charts demonstrate the fraction (%) of DEPs upregulated (red) or downregulated (blue), with a threshold of log_2_(FC) >0.8 or <-0.8 in Ctrl and KO mice (bottom). The gray part represents the fraction of DEPs with −0.8≤log_2_(FC<u>)<</u>0.8. n=3 per group. **D.** Top 15 pathways most enriched in upregulated (red) and downregulated (blue) DEPs in the lungs of Ctrl (left) and KO mouse (right) following MMC treatment. The green, red, and orange asterisks indicate pathways related to protein synthesis/modifications, RNA metabolism, and mitochondrial ATP synthesis, respectively. **E.** The bar graphs indicate the fold change of ribosomal proteins (top) and components of the mitochondrial electron transport chain (ETC) (bottom) between vehicle and MMC treatment in Ctrl and KO mice. Red and blue colors indicate upregulated and downregulated proteins. Darker colored bars represent *p* <0.05.

### PKR-depleted vascular endothelial cells are not impaired by MMC treatment

We previously demonstrated that MMC treatment induces the depletion of the adherens junction protein VE-Cadherin (VE-Cad, also known as Cdh5), resulting in increased vascular permeability (16, 17). To investigate this mechanism further, we isolated pulmonary vascular endothelial cells (PVECs) from the lungs of Ctrl and KO mice treated with either Veh or MMC. These cells were subjected to immunofluorescence (IF) staining for VE-Cad. PVECs from MMC-treated Ctrl mice exhibited a reduction in VE-Cad levels. In contrast, PVECs from MMC-treated KO mice retained VE-Cad expression at levels comparable to those observed in Veh-treated Ctrl or KO mice (**Fig. 3A**). Additionally, IF staining for p-PKR and ATF4 confirmed the activation of the ISR in PVECs from MMC-treated Ctrl mice, as evidenced by the induction of p-PKR in the cytoplasm and ATF4 in the nucleus (**Fig. 3B**). In contrast, PVECs from MMC-treated KO mice showed neither p-PKR staining nor ATF4 accumulation, indicating the lack of ISR activation (**Fig. 3B**). These results align with the immunoblot results obtained from lung lysates (**Fig. 1A**) and support that the activation of PKR-ATF4-ISR axis following MMC treatment occurs in vascular endothelium. Moreover, these results demonstrate that MMC does not mediate the downregulation of VE-Cad and impairs the endothelial barrier when PKR is ablated; therefore, ISR activation is prevented in the vascular endothelium.

We previously demonstrated that the VE-Cad and Rad51 Complex (VRC), located at the adherence junctions in PVECs, is depleted due to the release of VRC into circulation following MMC treatment in rats (16, 17). By immunoprecipitation with an anti-Rad51 antibody, followed by the immunoblot with an anti-VE-Cad antibody, we found a 2.1-fold increase in the amount of VRC in the plasma of Ctrl mice upon MMC treatment (**Fig. 3C**). Similarly to VRC, the levels of VE-Cad and Rad51 in the input plasma samples of Ctrl mice increased by 2.8-fold and 3.6-fold, respectively, following MMC treatment (**Fig. 3C**). In contrast, KO mice showed no induction of plasma VRC, VE-Cad, or Rad51 after MMC treatment (**Fig. 3C**). These results demonstrate that the junctional structure of the pulmonary vascular endothelium in KO mice is protected from MMC-mediated damage because of the ablation of PKR-mediated ISR activation and the release of VRC.

### Proteomic analysis reveals protection in KO Mice from MMC-mediated translational inhibition

One of the consequences of ISR activation is the attenuation of global cap-dependent translation due to the eIF2α phosphorylation (8, 9). To examine the effect of PKR-mediated ISR activation on the protein dynamics upon MMC treatment, we applied quantitative mass spectrometry (MS) analysis and compared the lung proteome between Veh- and MMC-treated Ctrl and KO mice. Principal component analysis (PCA) (**Fig. 4A**) revealed that the proteome of Veh-(Ctrl-Veh) and MMC-treated Ctrl mice (Ctrl-MMC) were highly distinct (**Fig. 4A**). In contrast, the proteome of Veh-treated (KO-Veh) and MMC-treated KO mice (KO-MMC) were more closely associated (**Fig. 4A**). Among the 7,598 proteins detected in the lungs of Ctrl and KO mice, 4,385 proteins in Ctrl mice and 1,583 proteins in KO mice were identified as differentially expressed, based on a *P*-value cutoff of 0.05 and a log_2_(FC) threshold of >0.8 or <-0.8. A hierarchically clustered heat map of differentially expressed proteins (DEPs) illustrated alterations in protein levels following MMC treatment and genetic ablation of PKR (**Fig. 4B**). In Ctrl mice, 68.6% of DEPs (3,006 proteins) exhibited decreased abundance after MMC treatment, whereas only 9.2% of DEPs (145 proteins) showed a decrease in KO mice (**Fig. 4C**). These observations support our results that, in KO mice, ISR activation and global translational attenuation were not triggered by MMC (**Fig. 1**). In contrast, 0.4% (17 proteins) and 14.3% (226 proteins) of DEPs showed increases in Ctrl and KO mice, respectively, following MMC treatment (**Fig. 4C**). Gene ontology (GO) enrichment analysis revealed five most overrepresented GO terms in DEPs downregulated in Ctrl mice were protein metabolism, including protein synthesis, modification, and localization (**Fig. 4D**, green asterisk). We also noted that proteins associated with RNA metabolism were overrepresented in downregulated DEPs in Ctrl mice (**Fig. 4D**, red asterisk). In KO mice, however, the GO terms overrepresented in downregulated DEPs in KO mice were divergent from those in Ctrl mice, such as platelet activation and the metabolic process of fatty acid, arachidonic acid, and vitamin D (**Fig. 4D**, bottom right). Indeed, 67 out of 73 ribosomal proteins (RPs) (92%) were downregulated in Ctrl mice following MMC treatment, resulting from the attenuation of protein synthesis through ISR activation by PKR (**Fig. 4E, top**). In contrast, only 21 out of 73 RPs (29%) were downregulated in KO mice, presumably due to a lack of PKR-ISR activation (**Fig. 4E, top**). We noted that the GO terms related to the mitochondrial electron transport chain (ETC) were over-represented in DEPs upregulated in MMC-treated Ctrl mice (**Fig. 4D**, orange asterisks). However, we found that 89% (56 out of 63) of the primary components of Complexes I–V in the ETC decreased following MMC treatment in Ctrl mice (**Fig. 4E, bottom**), which could result in a reduced ATP synthesis and an accumulation of reactive oxygen species(27). In contrast, in KO mice, 57% (36 out of 63) of these proteins were upregulated following MMC treatment (**Fig. 4E, bottom**). These results suggest that the MMC-mediated activation of the PKR-ISR axis results in robust proteomics changes, disrupts cellular homeostasis, and mediates vascular remodeling in PVOD. Moreover, genetic ablation of PKR protects against MMC-induced ISR activation, attenuation of protein synthesis, and PVOD pathogenesis. Thus, we identify PKR as a promising therapeutic target for PVOD.

## Discussion

In this study, we established a murine model of PVOD by administering MMC at the same dosage previously used to generate PVOD in Sprague-Dawley rats (16, 17). Like in rats, MMC-treated mice developed obliterative fibrosis in arterioles and venules, adventitial growth, and capillary hemangiomatosis, characteristics of PVOD patients (16, 17). Unlike in rats, where vascular remodeling becomes evident by day 8 and progresses to more severe manifestations by day 24, and the phenotypes remain the same up to day 60 with no mortality following MMC treatment (16, 17), Ctrl mice exhibited severe PVOD phenotypes, including pleural and pericardial effusion, by day 5. By day 14, all MMC-treated Ctrl mice died of severe pulmonary vascular remodeling in PAs and PVs. In contrast, no mortality was observed among MMC-treated KO mice, indicating the resilience of KO mice against the insult by MMC.

The biallelic mutation in the *Eif2ak4* gene, which encodes GCN2, is the primary genetic cause of PVOD(6). Our findings of the involvement of PKR activation in PVOD pathogenesis are seemingly at odds with the genetic observations that PVOD-associated mutations in *Eif2ak4* are believed to result in loss-of-function/expression mutations in GCN2, given that GCN2 and PKR share the same function(6, 16, 17). Recent studies on missense GCN2 mutants associated with PVOD show that some mutants are misfolded and degraded or kinase-inactive, while others are hypomorphic (28). Thus, it remains unclear whether all PVOD patients carrying *Eif2ak4* mutations develop the disease due to the inactivation of GCN2. It is plausible that genetic inactivation of GCN2 could lead to compensatory activation of PKR (8). The mechanisms driving the compensatory activation of PKR should be explored.

We showed previously that the inhibition of PKR by C16 prevents and reverses MMC-induced PVOD phenotypes in rats (16, 17). The observations in PKR-null mice validate that the reversal of PVOD phenotypes by C16 is indeed through the inactivation of the PKR-ISR axis and shed light on the essential role of PKR in MMC-induced PVOD pathogenesis in rodents. PKR was initially identified as a viral infection sensor that inhibits viral replication by suppressing host cell translation (19). In recent studies, however, PKR has been recognized to play a critical role in maintaining tissue homeostasis under sterile conditions (19). Mutations in PKR or its activator protein PACT, which result in increased PKR activation, are linked to the neurological disorder dystonia (29–31). Aberrant PKR activation has also been linked to autoimmune diseases, such as rheumatoid arthritis and systemic lupus erythematosus (19). Given the ubiquitous expression of PKR, examining the phenotypes of vascular endothelial cell-specific PKR knockout mice might provide further insight into the role of the PKR-ISR pathway, specifically in the endothelium.

We also provided evidence of the role of the PKR-ISR axis on the reprogramming of the proteome in vascular endothelial cells. It is well accepted that PKR, along with the other three eIF2 kinases, phosphorylates eIF2α and attenuates global cap-dependent translation as a critical component of the ISR (8, 9). The transient activation of the ISR and alterations of the proteostasis under stress are thought to enhance cellular resilience and facilitate adaptation to stress conditions (8, 9). However, the quantitative and qualitative changes in the proteome upon specific stress within a specific tissue have not been well characterized. Our proteomic analysis in Ctrl mice demonstrates that MMC treatment leads about a quarter of lung proteins to up or downregulate more than 1.74-fold. Among the DEPs downregulated in Ctrl mice, proteins associated with protein and RNA metabolism were significantly enriched following MMC treatment. MMC promotes constitutive eIF2α phosphorylation and translational inhibition in the pulmonary vascular endothelium of PVOD rats and patients through downregulating protein phosphatase 1 (PP1) complex that dephosphorylates Ser51 of eIF2α where PKR phosphorylates upon MMC treatment (16, 17). We speculate that constitutive eIF2α phosphorylation and ISR activation, as opposed to transient ISR activation, lead to the irreversible reprogramming of proteostasis and drive the deregulation of endothelial cells and pathologic remodeling of pulmonary vessels in PVOD. Future studies focusing on the pathways enriched in DEPs in Ctrl mice (Veh vs. MMC) will reveal previously unappreciated mechanisms underlying vascular remodeling in PVOD.

A total of 68.6% of the proteins reduced in Ctrl mice following MMC treatment, as identified by proteomic analysis, is close to the observations in the puromycin incorporation assay, demonstrating a 74% reduction in nascent proteins. The proteins with a long half-life may not appear to be substantially diminished after MMC treatment. This could lead to an underestimation of the impact of global translational attenuation induced by the PKR-ISR axis in our proteomics analysis. Stable isotope labeling by amino acids in cell culture (SILAC)-based proteomic analysis and ribosome profiling might better represent proteome dynamics following ISR activation (32, 33). It is well-established that cellular energy production and consumption are regulated by various environmental and physiological stressors (34, 35). Our results indicate that, following MMC treatment, most mitochondrial ETC components are downregulated in Ctrl mice. These findings suggest that PKR-dependent ISR activation reduces energy production by depleting ETC components while reducing energy consumption by suppressing RNA and protein metabolism. In conclusion, this study highlights the essential role of PKR in ISR activation and its impact on proteostasis, ultimately contributing to PVOD pathogenesis. Our findings suggest PKR inhibitors are promising candidates for therapeutic intervention in PVOD.

## Methods

Reagents, kits, antibodies, PCR primers, siRNAs, instruments, and software used in the study are listed in the Supplementary Information.

### Sex as a biological variable

Sex was not considered a biological variable in this study. Both male and female animals were used in all experiments.

### Study Approval

All animal experiments were conducted by the guidelines of the Institutional Animal Care and Use Committee (IACUC) at the University of California, San Francisco. The relevant protocol, AN200674-00B, was approved by the IACUC on October 08, 2024.

### Mouse model of PVOD

The generation of *EIF2AK2* knockout (KO) mice was previously described (21). Ctrl and KO mice in the C57Bl/6 background were housed in the vivarium of the cardiovascular research building at UCSF (San Francisco, USA). Both male and female young mice (9-10 weeks) were subjected to the following protocols to examine the effect of MMC. Two mg of MMC was dissolved in 1 ml phosphate-buffered saline (PBS). Mice were randomly divided into MMC (3 mg/kg body weight)- or vehicle (PBS)-exposed groups. MMC or vehicle was administered once intraperitoneal injection (i.p.) on day 0. Mice were euthanized on day 5 for hemodynamic measurements, RV hypertrophy assessment, and tissue collections.

### Hemodynamic assessment

To assess hemodynamics, terminal right heart catheterization was performed in mice using an open-chest method, as previously described (36, 37). Mice were anesthetized intraperitoneally with a mixture of ketamine (100 mg/kg) and xylazine (40 mg/kg). A tracheostomy tube was placed, and mechanical ventilation was initiated at a tidal volume of 6 cc/kg. Following anesthesia, the abdomen and diaphragm were carefully dissected to minimize blood loss. A 1 Fr pressure-volume catheter (PVR-1035, Millar AD Instruments, Houston, TX) was inserted directly into the right ventricle (RV) and subsequently into the left ventricle (LV) through their respective free walls. At the end of the hemodynamic measurements, the lungs were perfused with PBS via the RV. The Fulton index, calculated as the weight ratio of the right ventricle to the combined weight of the left ventricle and septum [RV/LV+S)], was used to measure RV hypertrophy. The left lung lobes were inflated with 1% agarose and fixed in 10% formalin for histological analysis, while the four right lung lobes were snap-frozen at −80°C for subsequent protein and RNA analysis.

### Histological assessment of vascular remodeling

Anesthetized mice were flushed with 1X PBS, fixed in 4% paraformaldehyde (w/v), transferred to 1X PBS after 24 h, and subjected to paraffin embedding. The right bronchus of flushed lungs was sutured, the left lung inflated with 1% low melt agarose for fixation and paraffin embedding, and the right lung split for snap freezing for protein and RNA studies.

To assess pulmonary artery and vein muscularization, mice lung tissue sections (10 µm in thickness) were subjected to conventional H&E staining. The external and internal diameter of a minimum of 50 transversally cut vessels in tissue block ranging from 25 to 80 µm was measured by determining the distance between the lamina elastica externa and lumen in two perpendicular directions described previously (38). The vessels were subdivided based on their diameter (microvessels: <50 µm and medium-sized vessels: 50-80 µm), and the assessment of muscularization was performed using ImageJ in a blinded fashion by a single researcher to reduce operator variability, which was not aware of the group allocation of the samples being analyzed. The vessels with >50% luminal occlusion are considered moderately occluded, and those with <50% are identified as severely occluded. The absolute count of the moderately and severely occluded vessels is the percent fraction of all counted vessels. Images were acquired using Ts2 microscopes (Nikon) and Leica SPE confocal microscope. Airways (bronchi and bronchioles) follow a branching pattern that mirrors the tree-like structure of the lung’s lobes and segments, while veins have a more variable course as they drain blood back to the heart. Airways generally have thicker walls compared to veins. When cut in cross-section, they are more likely to maintain a round or oval shape, while veins may appear more collapsed or irregular. These characteristics were applied to distinguish veins from airways.

### Microfil-casting of vasculature

The procedure for casting the vessels with Microfil was described previously(39). Briefly, heparin (1000 UI/kg), as an anticoagulant, was injected intravenously 10 min before anesthesia. The mice were anesthetized using a ketamine/xylazine cocktail. After removing the anterior chest wall, a microperfusion tube was inserted and kept in the right ventricle via a needle (25 G) for perfusion with PBS, which drained from the left atrium. The lungs were perfused to clear all blood, as evidenced by turning the lung tissue white. During the perfusion of pulmonary arteries (PAs), a freshly dissolved Microfil polymer mixture (MV compound: MV diluent: MV agent = 5:5:1) was instilled via a 25 G needle inserted into the PA from the incision through the RV wall by manual injection. The Microfil mixture was gently infused into the PA under a dissecting microscope until it reached the terminal branches of PAs and stopped within 2–3 sec. The lungs were then kept at room temperature for approximately 90 min or overnight at 4 while covered with a wet paper towel to avoid desiccation of the lungs. At the end of the experiment, the dissected lungs and hearts were rinsed in PBS for 10–15 min at room temperature. They were then dehydrated in ethanol solutions (50%, 70%, 80%, 95%, and 100%; 2 h each). After dehydration, the lungs were put into a methyl salicylate (Sigma-Aldrich) solution. When the lungs became translucent, and the Microfil was visible, they were photographed using a Nikon SMZ800N stereomicroscope. The number of branches and junctions in the distal pulmonary vascular networks of all lobes was counted by ImageJ.

### Immunoblot analysis

The mice tissue lysates were prepared in the lysis buffer (1% Triton X-100, 150mM NaCl, 50mM Tris-Cl at pH 7.5, 1mM EDTA). The supernatants were collected, and total protein concentration was measured by NanoDrop 2000c (Thermo Scientific). Protein samples were denatured in SDS-sample buffer for 5 min at 95°C, loaded onto Mini-Protean TGXTM gels (BioRad Laboratories) in equal amounts, and subjected to electrophoresis. Nitrocellulose membrane (Genesee Scientific) was used to blot the gels, which were blocked with 5% non-fat milk or 3% BSA in 1x Tris-buffered saline with 0.1% tween-20 (1x TBST) for one hour at RT. The membranes were incubated at four °C overnight with a primary antibody.

Chemiluminescence signals were detected using SuperSignal™ West Dura extended duration substrate (ThermoFisher) and imaged using an Odyssey Dlx Imaging System (LI-COR). Antibodies used for immunoblots are found in Supplementary Information. The quantity of each protein was normalized to the amount of a loading control protein. Subsequently, its relative quantity was calculated by setting the amount of the protein in the control (vehicle-treated) sample as 1.

### Immunoprecipitation assay

mice plasma samples were lysed in IP buffer (1% Triton X-100, 150 mM NaCl, 50 mM Tris-Cl at pH 7.5, 1mM EDTA) supplemented with protease inhibitors (1:100 dilution) and phosphatase inhibitor (1:100 dilution). Lysates were nutated for 30 min at four °C, followed by centrifugation at 12,000 g for 10 min, and supernatants were collected. One-tenth of the lysate was saved as an input sample for immunoblot. The lysate was incubated with indicated antibodies and anti-IgG (negative control) nutating overnight at 4°C, followed by the addition of Dynabeads^TM^ Protein A/G and rocking for four hours at four °C. The dynabeads were precipitated and rinsed thrice with IP buffer for 5 min at 4 °C. The eluate was boiled at 95°C for 8 min in a loading buffer and subjected to immunoblot along with input samples. For the input samples, the quantity of the indicated protein was initially normalized to the amount of a loading control protein, such as β-actin. Subsequently, its relative quantity was calculated by setting the amount of the protein in the vehicle-treated sample as 1. For the IP samples, the amount of the indicated protein in the MMC-treated sample was presented with the protein amount in the vehicle-treated sample set as 1.

### Reverse Transcriptase-quantitative Polymerase Chain Reaction (RT-qPCR)

Total RNA was extracted from mouse lungs and subjected to cDNA preparation by the reverse transcription reaction using an iScript cDNA Synthesis Kit (#17088890, Bio-Rad). qPCR analysis was performed in triplicate using iQ SYBR Green Supermix (#1708882, Bio-Rad). The relative expression values were determined by normalization to *GAPDH* transcript levels and calculated using the ΔΔCT method. RT-qPCR primer sequences are found in Supplementary Information.

### Chromatin immunoprecipitation (ChIP) assay

The lungs of mice treated with vehicle (saline) or MMC (3 mg/kg) were harvested on day 5. Lung cells were isolated and crosslinked with 1% formaldehyde for 15 minutes at room temperature (RT), followed by quenching with 1 M glycine. After washing the cells with 1x PBS, they were lysed using a lysis buffer containing 50 mM Tris-HCl (pH 8.1), 10 mM EDTA, 1% SDS, and a protease inhibitor. Genomic DNA was sheared to an average length of 200–500 bp by sonication, and the lysates were clarified by centrifugation at 12,000g for 10 minutes at 4°C. The supernatant was incubated with protein A/G Dynabeads at 4°C for 1 h, diluted the pre-cleared sample was to a 1:10 ratio with dilution buffer (20 mM Tris-Cl pH 8.1, 150 mM NaCl, 2 mM EDTA, 1% Triton X-100 and protease inhibitor) and 1/10 volume was kept as input before incubation with non-specific IgG (control), or an anti-ATF4 antibody overnight at 4°C followed by incubation with protein A/G Dynabeads. Next, the Dynabeads were washed with a buffer I (20 mM Tris-Cl pH 8.1, 150 mM NaCl, 2 mM EDTA, 1% Triton X-100, 0.1% SDS), buffer II (20 mM Tris-Cl pH 8.1, 500 mM NaCl, 2 mM EDTA, 1% Triton X-100, 0.1% SDS), and buffer III (10mM Tris-Cl pH8.1, 250mM LiCl, 1mM EDTA, 1%NP-40, 1% Deoxycholate) at 4°C. The Dynabeads were washed twice with cold TE (10 mM Tris-Cl pH 8.1, 1 mM EDTA) and incubated in 250 μl elution buffer (200 mM NaHCO_3_, 1% SDS) at RT for 15 min. The eluates were mixed with 1/25 volume 5M NaCl and incubated at 65 °C for 4 h. 1/50 volume of 0.5 M EDTA, 1/25 volume of Tris-Cl pH 6.5, proteinase K (final 100 μg/ml) were added and incubated at 45°C for 1 h. Immunoprecipitated DNA fragments were purified with a QIAquick PCR Purification Kit, followed by RT-qPCR analysis. The sequences of PCR primer for ChIP assay are found in Supplementary Information.

### Isolation of pulmonary vascular endothelial cells

CD31-positive pulmonary vascular endothelial cells (PVECs) were isolated from the lung tissues by enzymatically digestion method as previously described (40). Digestions were stopped after 20 min, and cells were processed into a single-cell suspension on ice. Cells were sorted with CD31 magnetic beads and grown on the coverslips for immunofluorescence staining.

### Immunofluorescence staining

The endothelial cells isolated from mice lung vasculature were blocked and incubated with primary antibody overnight at 4°C. Alexa Fluor secondary antibodies (Invitrogen) were applied for 2 hours at RT. IF images were acquired using a confocal microscope (Leica SPE) or Eclipse Ts2 Inverted LED phase contrast microscope (Nikon) and analyzed using ImageJ. Antibodies used are found in the Supplementary Information.

### In vivo puromycin incorporation assay

Mice were injected once with either a vehicle (saline) or MMC (3 mg/kg). Five days after the injection, the mice were perfused with 100 µg/ml puromycin in 1XPBS via intracardiac injection. Ten minutes following the perfusion, the lungs were harvested, and total lung lysates were subjected to SDS-PAGE. Immunoblot was conducted using an anti-puromycin antibody.

### Mass spectrometry sample preparation

The mice lung samples (50 mg) were lysed with an 8M Urea buffer supplemented with a 1X protease inhibitor cocktail (Thermo Fisher Scientific, 1861280). Cells were then homogenized via sonication, followed by a strong spin at 17,000xg for 10 minutes at 4°C to clear the debris and extract the protein-containing supernatant. Protein concentration was measured using a BCA kit per the manufacturer’s instructions. 100µg proteins were taken for further processing and resuspended in a digestion buffer (PreOmics, P.O.00027), followed by the addition of trypsin for in-solution peptide digestion. The digestion process was performed for 90 minutes at 37°C while shaking at 700 rpm. The resulting peptide mixture was then desalted, eluted, and dried completely using a vacuum concentrator (Labconco, 7810010). Dried peptides were reconstituted in 2% acetonitrile (ACN) and 0.1% formic acid, and their concentration was measured on a NanoDrop using Protein205A and 280A (Thermo Fisher Scientific). Finally, the peptide concentration was adjusted to 0.1 μg/μL for mass spectrometry analysis.

### Liquid chromatography and Mass spectrometry analysis

Mass spectrometry analysis was performed using a timsTOF Pro2 instrument (Bruker Daltonics) coupled to a nanoElute UHPLC system. Peptides were separated on a 150 µm × 25 cm C18 column (1.5 µm particle size, Bruker) at 50°C, with mobile phases comprising 0.1% formic acid in water (solvent A) and 0.1% formic acid in acetonitrile (solvent B). A 60-minute gradient was employed at a flow rate of 300 nL/min: 2% solvent B for 5 minutes, followed by a linear increase to 30% solvent B over 60 minutes, followed by column equilibration and washing. The timsTOF Pro2 was operated in parallel accumulation-serial fragmentation (DIA-PASEF) mode using data-independent acquisition settings. The MS scan range was considered from 100-1700 m/z in positive polarity with isolation widths of 25 Da. The TIMS setting was kept in custom ion mobility mode with 1/K0 starting at 0.7 V.s/cm^2^ to end at 1.3 V.s/cm^2^, with a frame cycle of 1.2 s, incorporating 120 ms ion accumulation per frame. On average, 12 PASEF scans were acquired per duty cycle. The raw LFQ spectral spectra were analyzed using DIA-NN, a neural network-based software, to identify and quantify surface proteins (PMID: 31768060). The data were searched against the Uniprot-reviewed mouse proteome database. Trypsin was specified as the protease, allowing for up to one missed cleavage. Methionine oxidation and N-terminal acetylation were set as variable modifications, while cysteine carbamidomethylation was fixed. The analysis was conducted in library-free mode, with DIA-NN generating in silico spectral libraries directly from the protein sequence database. Default settings for precursor charge states and ion types were used for protein quantification. The LFQ intensities were median normalized and subsequently log-transformed. Differential expression of proteins was assessed using Welch’s t-test, with significance defined by a p-value < 0.05 and a fold change (FC) threshold of greater than 1.74 or less than −1.74 to identify upregulated or downregulated proteins (41).

### Mass spectrometry data analysis

Raw data files were processed with Byonic software (Protein Metrics, Cupertino, California). Fixed modifications included +113.084 C. Variable modifications included Acetyl +42.010565 N-term, pyro-Glu −17.026549 N-term Q, pyro-Glu −18.010565 N-term E. Precursor tolerance 30.0 ppm.

### Data compilation

Raw files were read for UniProtIDs, gene names, and their respective mappings. ‘nan,’‘’ (empty strings), and ‘2 SV’ were ignored. Mappings between gene names and UniProtIDs were not one-to-one. Some genes and UniProtIDs were unmapped. Therefore, a comprehensive, one-to-one mapping was first made before compiling the raw data. Using the Retrieve/ID mapping program at www.uniprot.org (release 2019–11), all gene names were mapped to all possible UniProtIDs, and UniprotIDs were mapped to all possible gene names. The mappings from raw files and UniProt were combined to group equivalent gene names and UniProtIDs (usually with isoforms) together. From each group, one gene name and one UniProtID were selected for downstream data compilation. Any UniProtID or gene name that was not mapped was either given a protein ID (UNM #) or a gene name (Unm #). Thirty-eight gene names had no UniProtIDs, and 8 UniprotIDs had no gene names. All raw files were compiled into a single data table with the selected UniProtIDs and gene names (Supplementary Table 1).

### Statistical analysis

All numerical data are presented as mean ± standard error of the mean (SEM). Statistical analysis was performed using Microsoft Excel and GraphPad Prism 10 (La Jolla, CA). Datasets with two groups were subjected to a 2-tailed Student’s t-test, unpaired, equal variance, whereas comparison among three or more than three groups was made by ANOVA followed by Tukey’s post hoc corrections. Analysis of variance was applied to experiments with multiple parameters, one- or two-way, as appropriate. Significance was analyzed using a post hoc Tukey test and indicated as P-values where required. A *p*-value less than 0.05 was considered significant.

## Data availability

Raw proteomic data generated here will be deposited in the ProteomeXchange/PRIDE repository.

## Author Contributions

The authors confirm their contribution to the paper as follows. *Designing research studies*: A.P. and A.H. *Conducting experiments and acquiring data*: A.P. M.W., R.K., A.G., J.L., A.A., S.G., P.G. and A.B. *Analyzing data*: A.P. M.W., R.K., A.G., J.L., A.A., S.G., P.G., A.B., A.P.W., G.L., and A.H. *Writing the manuscript*: A.P., R.K., A.P.W., B.B.G., G.L., and A.H.

## Acknowledgments

We thank Drs. Hotamisligil (Harvard School of Public Health) and Biao Wang (UCSF) for sharing KO mice. Funding was provided by the National Heart, Lung, and Blood Institute (NHLBI; R01HL153915, and R01HL164581) to A.H.; American Heart Association 19CDA34730030, ATS Early Career Investigator Award in the Pulmonary Vascular Disease Program, and UCSF Resource Allocation Program grant to R.K.; by NHLBI (R01HL135872 and P01HL152961) to B.B.G; by NCI (R01CA290875) to A.P.W.

---

**Supple. Fig. 1.**
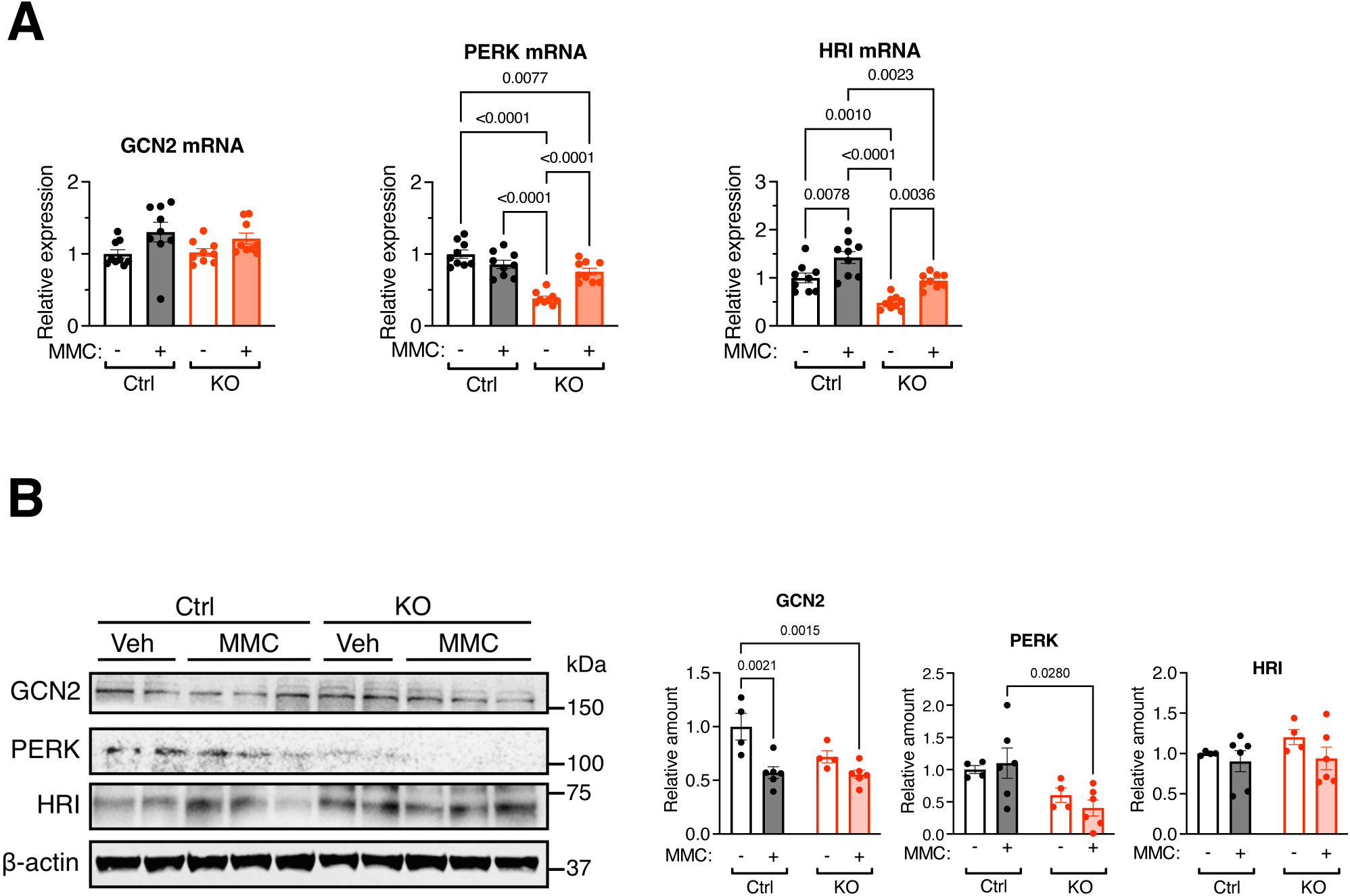
The levels of eIF2 kinases, such as GCN2, PERK, and HRI, are not altered in PKR KO mice. **A.** The level of mRNAs of GCN2, PERK, HRI target genes in the lung of from Ctrl and KO mice administered with vehicle or MMC was analyzed by qRT-PCR on day 5 and shown as mean±SEM. n=6 independent samples. **B.** The amount of GCN2, PERK, HRI, and β-actin are analyzed by immunoblots (left). The relative amounts of the indicated proteins, normalized to β-actin, are shown as mean ± SEM (right). *n* = 2-3 independent samples.

**Supple. Fig. 2.**
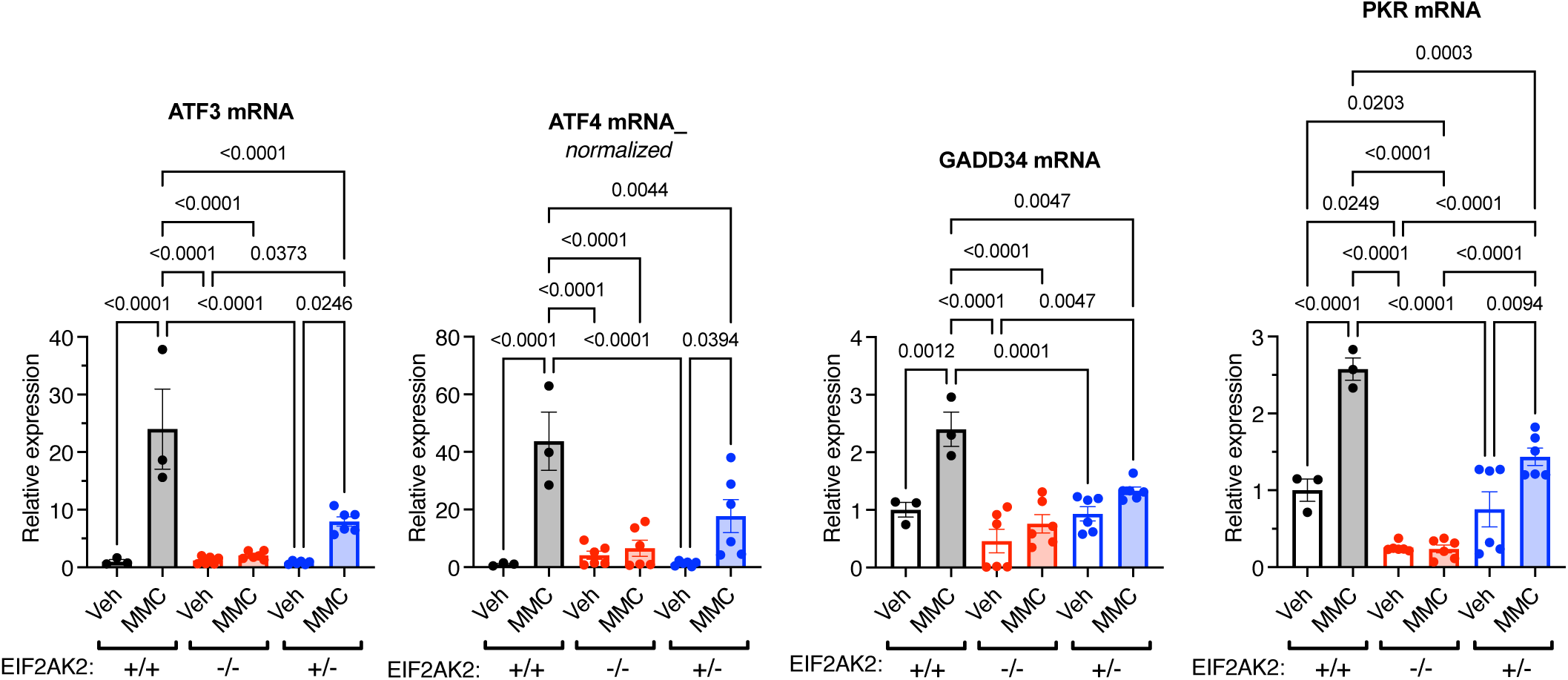
Partial induction of ATF4 target genes in the PKR heterozygous null mice. The level of mRNAs of ATF4 target genes, such as ATF3, ATF4, GADD34, and PKR in the lung of *EIF2AK2* wild type (+/+) mice (black)*, EIF2AK2* homozygous-null (−/−) mice (red), and *EIF2AK2* heterozygous-null (+/-) mice (blue) administered with vehicle or MMC was analyzed by qRT-PCR and shown as mean<u>+S</u>EM. n=6 independent samples.

**Supple. Fig. 3.**
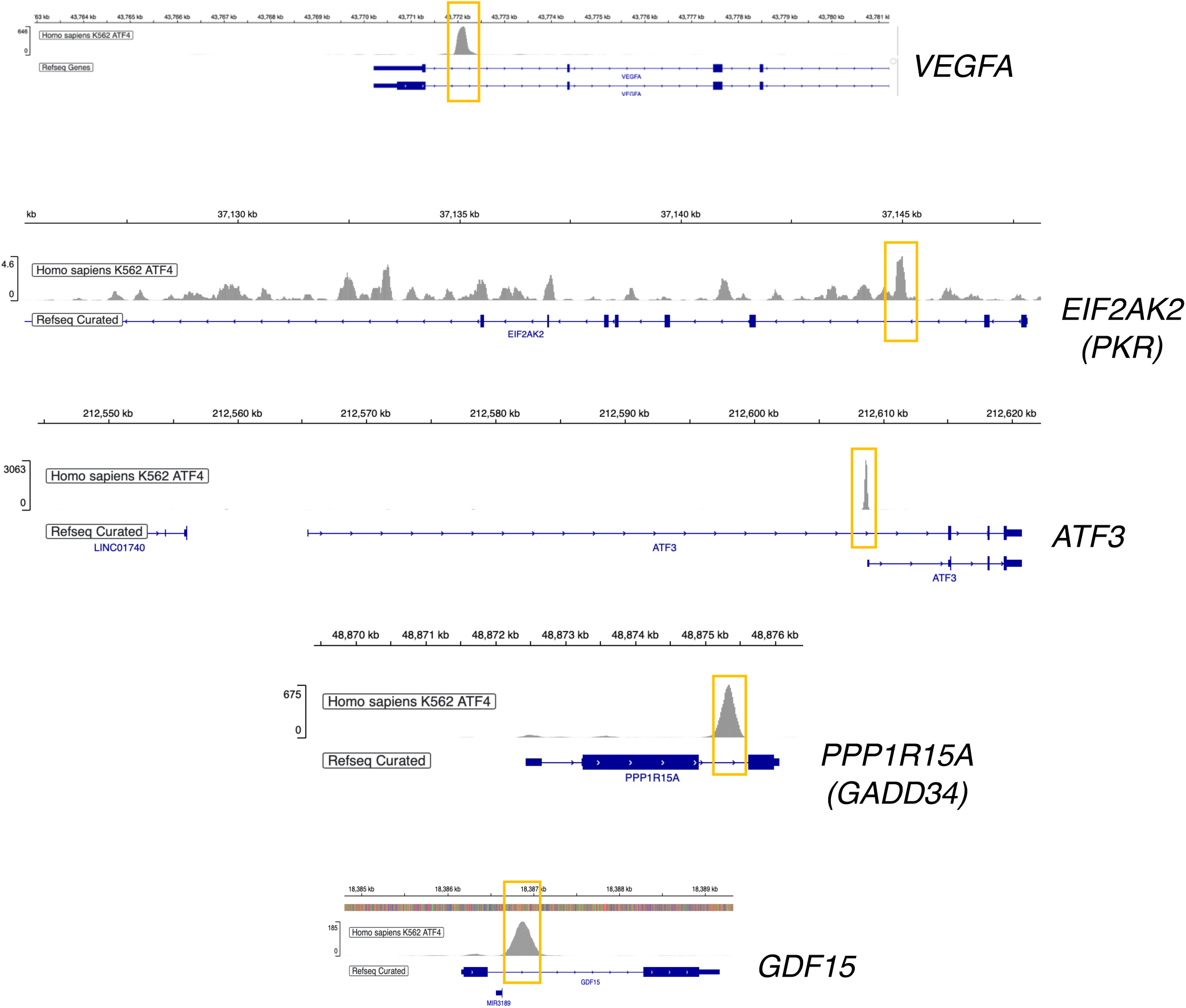
ChIP-seq maps the binding sites of ATF4 in the human genome. ChIP-seq data of the ENCODE database (Accession No. ENCFF484GNY and ENCFF742FPU) in human K562 cells identifies ATF4 binding sites in ATF4 target genes, such as *VEGFA*, *EIF2AK2*, *ATF3*, *PPP1R15A*, and *GDF15*. Orange rectangles indicate the genomic regions enriched by the ATF4 ChIP-seq.

**Supple. Fig. 4.**
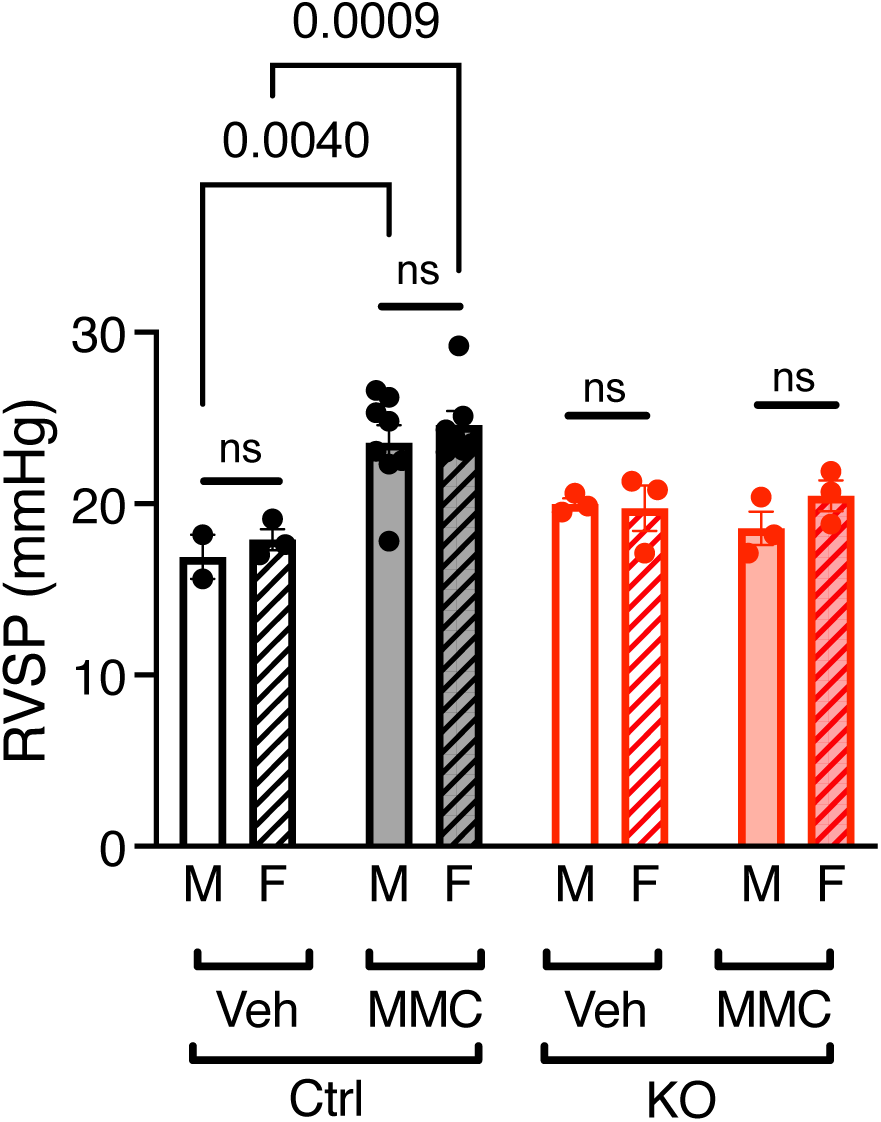
MMC-induced PVOD phenotype is comparable between male and female mice. RVSP (mmHg) was measured in the vehicle (Veh) or MMC-administered control (Ctrl) and KO mice and compared between male (M) and female (F) mice. n=2-8 per group.

**Supple. Fig. 5.**
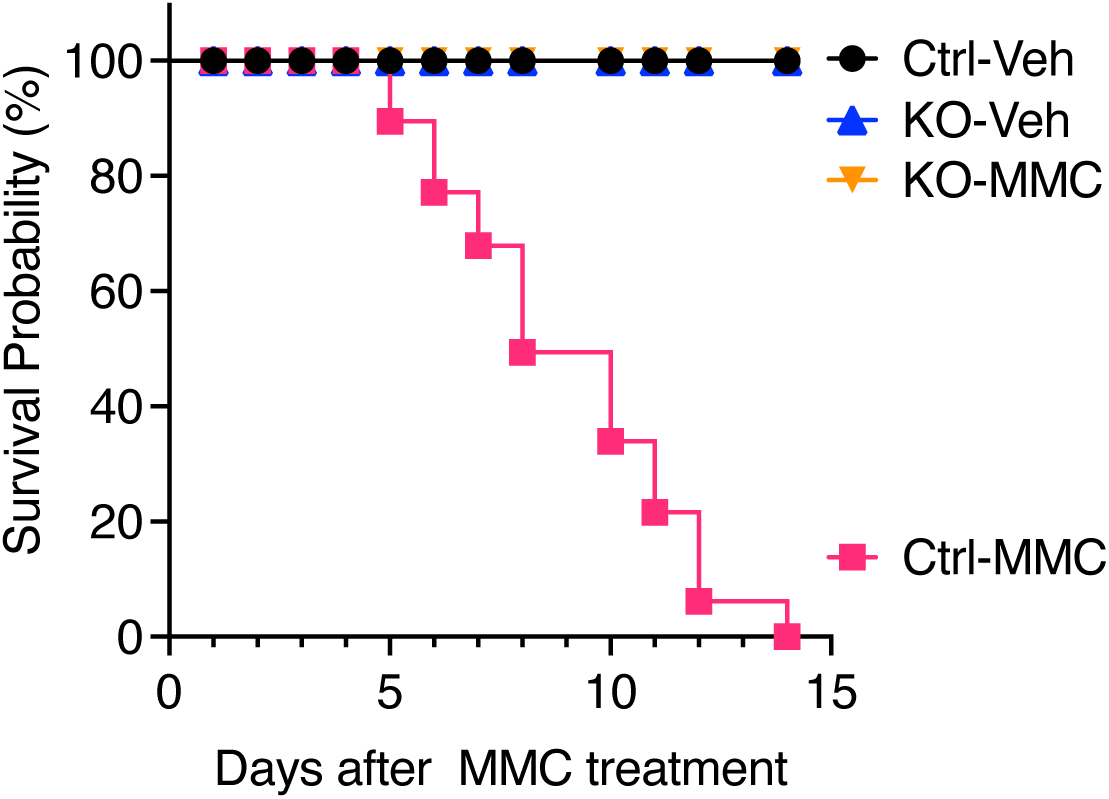
Kaplan Meier Survival curve of Ctrl and KO mice following MMC treatment. These cohorts include male and female mice of 9-10 weeks old. There is no mortality among Veh-treated Ctrl and Veh- or MMC-treated KO mice.

**Supple. Fig. 6.**
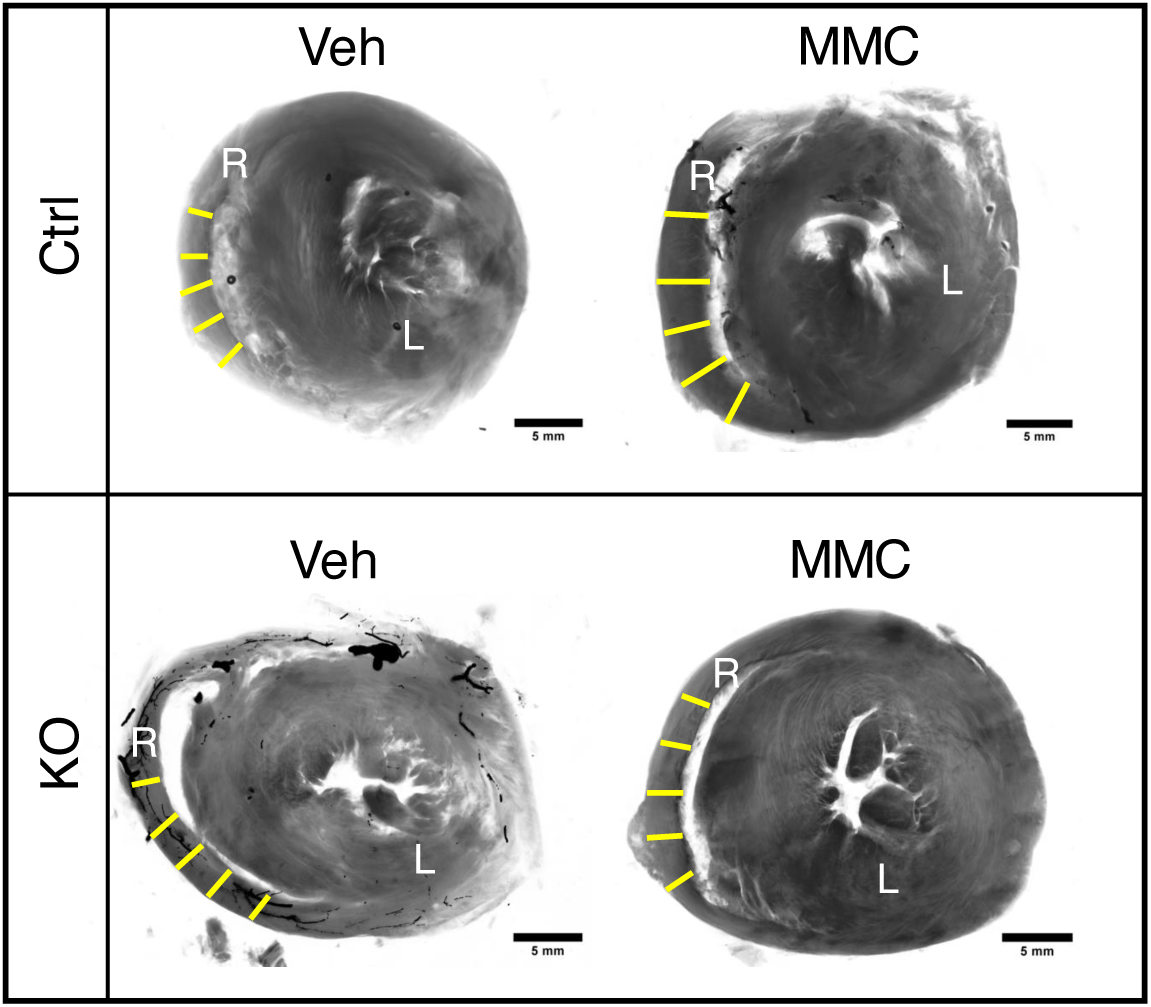
KO mice do not develop right ventricular hypertrophy following MMC treatment. Representative transverse sections of hearts isolated from Veh- or MMC-treated Ctrl and KO mice are shown. The right ventricle (R) and left ventricle (V) are indicated. Wall thickness was measured as depicted by the yellow lines, and the mean values were calculated to determine the right ventricular (RV) wall thickness, as presented in Fig. 2B. Scale bar=5 mm.

**Supple. Fig. 7.**
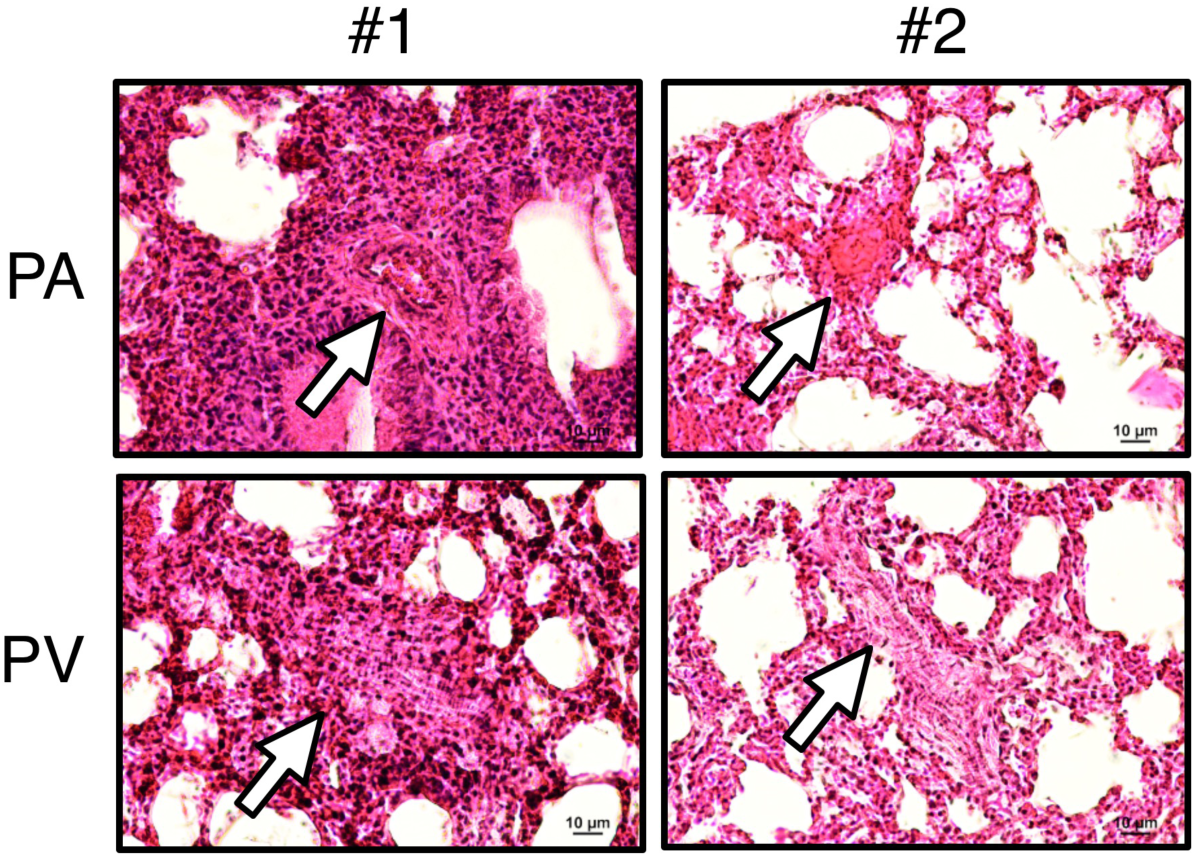
Fully occluded pulmonary vessels found in MMC-treated Ctrl mice that died after MMC administration. H&E staining images of PA and PV in two Ctrl mice (#1 and #2) that died on day 5 following administration of MMC are shown. White arrows indicate vessels. Scale bar=10 μm.

**Supple. Fig. 8.**
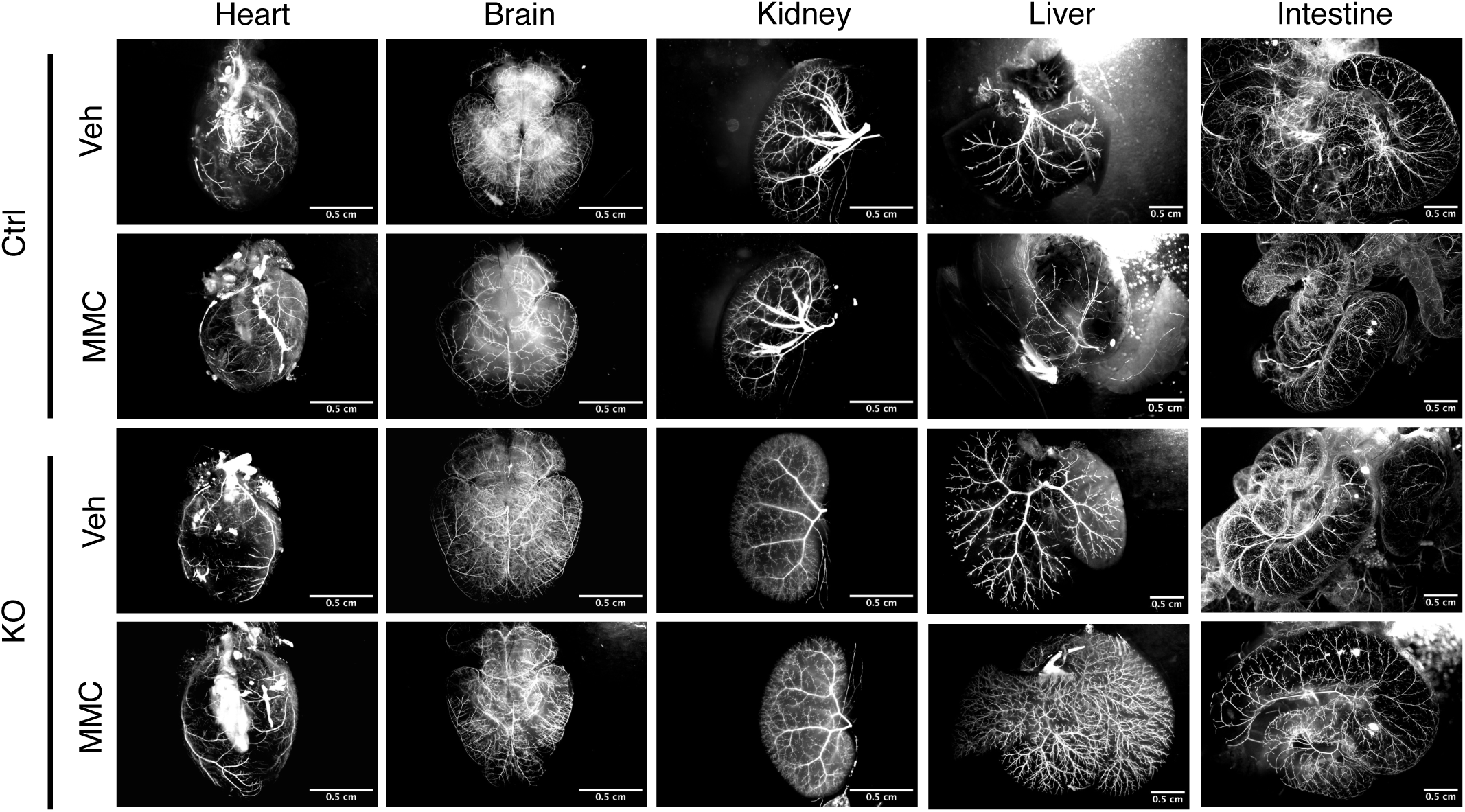
MMC-induced vascular remodeling is restricted within lung. Microfil casting of the vasculature in the heart, brain, kidney, liver, and intestine of Ctrl and KO mice treated with either Veh or MMC on day 5. Holistic images of the entire lung are displayed on the left, with a scale bar representing 0.5 cm. The number of branches and junctions per cm² of distal pulmonary vessels was quantified, with the data presented as mean ± SEM (right). n = 3 independent samples per group.

